# EPI-589, a redox-active neuroprotectant, potently protects cultured cells from oxidative stress and alleviates symptomatic and pathological progression of motor neuron disease in the wobbler mouse

**DOI:** 10.1101/2022.03.13.484182

**Authors:** Yuji Matsumoto, Kazuaki Sampei, Tetsuaki Nashida, Yuta Fujii, Naoko Tani, Fumiaki Ishibashi, Mitsugu Yamanaka, Takeo Ishiyama

**Author notes:** **Corresponding author** Yuji Matsumoto, Drug Research Division, Sumitomo Dainippon Pharma Co., Ltd., 6-8, Doshomachi 2-chome, Chuo-ku, Osaka, Osaka 541-0045, Japan. **Author e-mail addresses** Yuji Matsumoto Kazuaki Sampei, Tetsuaki Nashida, Yuta Fujii, Naoko Tani, Fumiaki Ishibashi, Mitsugu Yamanaka, Takeo Ishiyama.

## Abstract

Oxidative stress is believed to play a significant role in the pathophysiology of amyotrophic lateral sclerosis (ALS), the most common form of motor neuron disease. The present study aims to firstly investigate the antioxidant activities of EPI-589, a small-molecule quinone derivative, under cell-free or cell culture conditions, and explore the in vivo efficacy of EPI-589 in the wobbler mouse model of human motor neuron disease. The reduced form of EPI-589 showed hydroxyl radical scavenging activities, whereas the oxidized form i.e. EPI-589 did not. In cellular models utilizing ALS patient-derived fibroblasts carrying mutations in the fused in sarcoma (FUS) gene or superoxide dismutase 1 (SOD1) gene, EPI-589 potently protected cells from oxidative stress induced by buthionine sulfoximine and ferric citrate. Protective effect of EPI-589 was also observed in culture of mouse immortalized striatal STHdHQ7/Q7 cells with cystine deprivation. In wobbler mice, oral administration of dietary EPI-589 provided long-lasting amelioration of both of deterioration of the rotarod walking performance and progression of forelimb deformity in wobbler mice throughout the treatment. In separate studies, we found that EPI-589 significantly suppressed changes of pathophysiological markers such as plasma phosphorylated neurofilament heavy chain, urinary 8-hydroxy-2’-deoxyguanosine, and cervical N-acetylaspartate in untreated wobbler mice. Thus, the present study firstly demonstrates that EPI-589 is a highly potent, redox-active neuroprotectant and robustly delays the symptomatic and pathophysiological progression of motor neuron disease in the wobbler mouse, and these findings strongly encourage further exploration of the therapeutic potential of EPI-589 for the treatment of ALS.

## 1. Introduction

Amyotrophic lateral sclerosis (ALS) is the most common form of motor neuron disease and has a cumulative life-time prevalence of about 1 in 400 individuals (Brown and Al-Chalabi, 2017). It is characterized by degeneration of motor neurons in the brain and spinal cord, relentlessly progressive paralysis of most skeletal, bulbar, and respiratory muscles, and typically, death within 3 to 5 years after clinical onset. Despite the devastating clinical features of ALS, only a few drugs are available, including riluzole and edaravone, which show modest efficacy to delay disease progression and thereby to improve survival or motor function (Miller et al., 2012; Edaravone (MCI-186) ALS 19 Study Group, 2017). Thus, there is an urgent need for more efficacious therapies.

Oxidative stress is believed to play a significant role in the pathophysiology of ALS (Barber and Shaw, 2010). A number of studies using various markers of the effect of oxidative stress on lipids, proteins, and DNA in postmortem neuronal tissues, cerebrospinal fluids (CSF), plasma, and urine provide robust evidence of increased oxidative stress in ALS patients (D’Amico et al., 2013). For example, using 8-hydroxy-2’-deoxyguanosine (8-OHdG), a marker of oxidative DNA damage, one study demonstrated a prominent increase in this marker in the ventral horn of the spinal cord in ALS patients (Ferrante et al., 1997), whereas another report observed an increase in the same marker in CSF, plasma, and urine in ALS patients vs. control subjects and a correlation of the rate of increase in the marker with disease severity (Bogdanov et al., 2000). From another point of view, dozens of causative gene mutations have been identified to date in familial and sporadic cases of ALS. These gene product functions imply that the major pathogenic pathways leading to ALS may involve mitochondrial dysfunction and oxidative stress (Cook and Petrucelli, 2019). Interestingly, the studies using cell and animal models with some of these ALS-linked gene mutations consistently revealed concomitant mitochondrial dysfunction and oxidative stress that eventually gave rise to motor neuron degeneration (Carrì et al., 2015; Lopez-Gonzalez et al., 2016; Wang et al., 2016; Moujalled et al., 2017). Furthermore, edaravone, an antioxidant against reactive oxygen species (ROS), has been shown to delay the progression of motor neuron degeneration in multiple animal models of ALS such as the superoxide dismutase 1 (SOD1) transgenic mouse (Ito et al., 2008) and wobbler mouse with a naturally occurring motor neuron disease (Ikeda and Iwasaki, 2015), and has been approved based on clinical trial results in patients with early stage ALS that demonstrate a 33% reduction in motor function decline with edaravone vs. placebo (Edaravone (MCI-186) ALS 19 Study Group, 2017). Therefore, oxidative stress appears to be an appropriate therapeutic target in both familial and sporadic ALS, and creating a more effective drug than edaravone to reduce oxidative stress may be a viable way to improve currently available therapy for this enigmatic disease.

The wobbler mouse has been extensively investigated as a naturally occurring mouse model of human motor neuron disease with a L967Q mutation of the gene encoding vacuolar protein sorting 54 (VPS54), a component of the Golgi-associated retrograde protein (GARP) complex (Schmitt-John et al., 2005). The wobbler mouse has an early clinical onset at 3 to 4 weeks of age, progressive muscle atrophy and paralysis predominantly in the forelimbs at 2 weeks of age (Mitsumoto and Bradley, 1982), and progressive loss of the motor neurons most extensively from 3 to 6 weeks of age (Pollin et al., 1990). Quantitative and semi-quantitative assessments of clinical symptoms, physiological analysis of muscular function, and histometric analysis of motor neurons and forelimb muscles have been developed and utilized in this animal model to evaluate a variety of investigational drugs for ALS, including neurotrophic factors (Mitsumoto et al., 1994; Mitsumoto et al., 2001), riluzole (Ishiyama et al., 2004), edaravone (Ikeda and Iwasaki, 2015), and others (Moser et al., 2013). The wobbler mouse is considered to be particularly useful for testing novel drugs targeting oxidative stress, because disease pathology in the wobbler mouse involves abnormality of the mitochondria (Santoro et al., 2004; Moser et al., 2013) as well as abnormality of the endoplasmic reticulum and massive activation of astrocytes and microglia in the cervical cord (Moser et al., 2013), all of which are known to be the major sources of reactive oxygen species, and the clinically approved antioxidant drug edaravone and others were found to be effective in delaying disease progression (Ikeda and Iwasaki, 2015; Moser et al., 2013). Comparisons of the efficacy of clinically approved drugs such as riluzole and edaravone are feasible in the wobbler mouse model, which is further considered to be suitable for predicting the appropriate clinically meaningful efficacious dose range of a new investigational drug.

EPI-589, known as (R)-troloxamide quinone, is a small-molecule quinone derivative (Fig. 1). We have preliminarily demonstrated that EPI-589 seemed to act as redox-active neuroprotectant under experimental conditions relevant to ALS (Motodate et al., 2019; Nashida et al., 2019). This present paper aims to firstly describe in detail the potent antioxidant activities of EPI-589 under various cell culture conditions and explore the *in vivo* efficacy of EPI-589 for motor neuron disease using the wobbler mouse model. The studies were designed to compare the *in vitro* and *in vivo* effects of EPI-589 with those of edaravone. For the study in the wobbler mouse to support in detail the protective effects of EPI-589 against motor neuron injury, we have newly examined the level of 8-OHdG in urine, pathological change in the Golgi apparatus of motor neurons with the Golgi marker MG-160 (Fujita and Okamoto, 2005), and the release into plasma of phosphorylated neurofilament heavy chain (pNfH) as a marker for neuronal degeneration (Li et al., 2016), as well as examined by magnetic resonance spectrometry (MRS), N-acetylaspartate level in the cervical cord as a marker for neuronal integrity (Carew et al., 2011). In addition, we have done quantitative and semi-quantitative assessments of clinical symptoms and histiometric analysis of the cervical cord motor neurons as previously described (Ishiyama et al., 2004). As a result, we fully report in this paper that EPI-589 is central nervous system-penetrable and more potent than edaravone in terms of its ability to protect cells against oxidative stress under various cell culture conditions, and it also exerts highly robust effects that delay motor neuron degeneration in the wobbler mouse. Hence, we provide useful information for further clinical development of EPI-589 for the treatment of ALS.

**Fig. 1.**
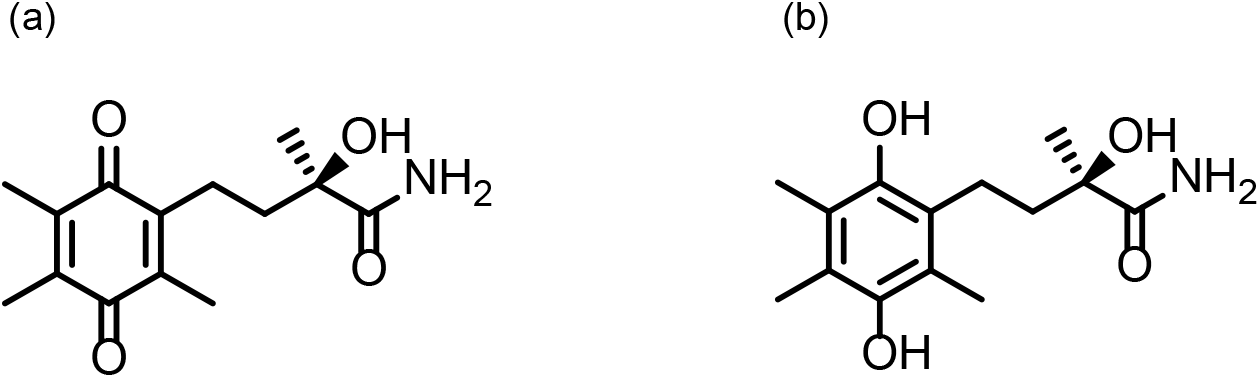
Chemical structure of EPI-589 (a) and the reduced form DSR-303359 (b).

## 2 Materials and Methods

### 2.1. Drugs

EPI-589 ((R)-2-hydroxy-2-methyl-4-(2,4,5-trimethyl-3,6-dioxocyclohexa-1,4-dien-1-yl)butanamide) and DSR-303359, a reduced form of EPI-589, were synthesized by Sumitomo Dainippon Pharma Co., Ltd. Edaravone hydrochloride (edaravone) was purchased from Sigma Aldrich (St. Louis, MO, USA). For the administration in rodent models, EPI-589 was mixed with powdered diet (CE-2; CLEA Japan, Inc., Tokyo, Japan) at a concentration of 10 mg/g using a mortar, and edaravone was dissolved in saline.

### 2.2. Animals

All experimental procedures for the use of animals were reviewed and approved by the Institutional Animal Care and Use Committee of Sumitomo Dainippon Pharma, Co., Ltd. Wobbler mice were originally obtained from the colony at the Cleveland Clinic Foundation Biological Resource Unit. The colony was expanded and maintained by KAC Co., Ltd. We obtained the mice as litters of 2-week-old pups with foster females. Homozygous affected wobbler mice were identified at 3 – 4 weeks of age by the appearance of the first clinical symptoms such as tremors and jitters. The accuracy of this clinical diagnosis has been established by histological studies (Mitsumoto and Bradley, 1982; Ishiyama et al., 2004), and reinforced by genotyping of the Vps54 gene mutation (Schmitt-John et al., 2005) conducted in another set of wobbler mice in our laboratory (data not shown). Immediately after the diagnosis, each wobbler mouse was transferred to a separate cage with a normal healthy female littermate as a foster mouse, and the experiments were started. All animals were kept in a room under controlled environmental conditions (temperature: 23 ± 3°C, humidity: 55 ± 15%, 12 hr light-dark cycle with light on at 8 am and used after a quarantine period of 7 days. The animals were given food and filtered tap water ad libitum.

### 2.3. Experimental design of wobbler mouse study

One behavioral study and two biomarker studies i.e. biomarker study 1 and biomarker study 2, were conducted using wobbler mice. In the behavioral study, motor function and clinical symptoms were repeatedly assessed in a blinded manner as described in detail below from 4 to 9 weeks of age, and then the plasma concentrations of compounds were measured at the end of treatment. Sixteen wobbler mice were initially assigned to either the control (no drug treatment), EPI-589, or edaravone group; however, 2 mice in the edaravone group were excluded from the analysis due to death, and 2 mice in control group and 3 mice in EPI-589 group were also excluded because of hydrocephalus as confirmed through brain autopsy after the sacrifice at the end of treatment.

Biomarker study 1 was conducted to evaluate plasma pNfH level in comparison with histopathological analysis of cervical cord motor neurons. The plasma sample for measurement of pNfH was collected from wobbler mice (n = 12) that were fed the control diet (CE-2) or dietary EPI-589 for 2 weeks after clinical symptom onset (from 4 to 6 weeks of age). At the end of treatment, four mice per group were arbitrarily chosen and sacrificed by transcardial perfusion with saline under isoflurane anesthesia, then the spinal cord was collected and fixed in 10% neutral buffered formalin for further immunohistological analysis. Biomarker study 2 was conducted to evaluate 8-OHdG level in urine and N-acetylaspartate (NAA) level in cervical spinal cord. Each urine sample for measurement of 8-OHdG was collected from wobbler mice (n = 12) fed the control diet or dietary EPI-589 for 3 weeks after clinical symptoms onset (from 4 to 7 weeks of age), whereas six mice per group were arbitrarily chosen to have continued drug treatment for another week by 8 weeks of age and then the cervical cord was obtained to evaluate NAA level. For these biomarker studies, age-matched, wild type littermate mice were also used to control for effects on homozygous wobbler mice. Blood samples were collected using heparinated tubes to obtain plasma for drug concentration and biomarker measurement.

### 2.4. Behavioral study

Prior to selection of mice for the study, all of the wobbler mice were acclimatized to the rotarod apparatus for 3 days, immediately after the diagnosis with clinical symptoms. Only the animals that walked for 300 seconds on the rotarod (10 rpm) during the acclimatization period were selected for use in this study. Then, the selected mice were allocated so as to ensure a gender balance in each group, and pre-administration values of rotarod performance, body weight, and forelimb deformity were confirmed to be comparable among the groups by Bartlett’s test.

Wobbler mice were administered dietary EPI-589 (10 mg/g of diet) or edaravone (15 mg/kg, i.p., daily) from 4 weeks of age, only after the clinical symptoms appeared. None of the wobbler mice in the EPI-589-treatment and control groups received intraperitoneal injections during the behavioral study. The edaravone dose in this study was same as that used in a previous report (Ikeda and Iwasaki, 2015). Rotarod walking time and forelimb deformity score were assessed once or twice a week in a blinded manner as briefly described below.

### 2.5. Rotarod performance test

In this study, the rotarod performance test in wobbler mice was done as previously reported (Ishiyama et al., 2004), by utilizing the MK-610A rotarod apparatus (Muromachi Kikai Co., Ltd., Tokyo, Japan). The animals were put on the rotating rod set at 10 rpm, and the time spent on the rod before falling was recorded with 300 seconds as the maximum. For each animal, measurement was repeated three times, and the maximum time was used for analysis.

### 2.6. Assessment of forelimb deformity score

The mouse forelimb deformity score is defined below with reference to the literature (Mitsumoto et al., 1994). Muscle atrophy and each forelimb deformity level were determined. Forelimb deformity score was the sum of both forelimb scores. Grade criteria: 0, normal; 1, atrophied paw (thin fingers); 1.5, over grade 1 but not grade 2; 2, curled digits; 2.5, over grade 2 but not grade 3; 3, curled wrist; 3.5, over grade 3 but not grade 4; 4, forelimb flexion contracture.

### 2.7. Measurement of drug concentration

In the preliminary study, four wobbler mice were fed dietary EPI-589 (10 mg/g of diet) and the plasma concentration of EPI-589 was measured as described below at 7 time points, every four hours for 24 hours after the start of the light cycle (8:00 AM). The study demonstrated that the mean plasma concentration of EPI-589 was maintained between 737 ± 112 and 2970 ± 2050 ng/mL as relatively lower and greater values in the light and dark cycle, respectively. At the end of treatment for EPI-589 in the behavioral study, the animals were sacrificed to measure drug concentration in plasma immediately after the start of the light cycle, and these drug concentrations were therefore supposed to be at the relatively lower end of maintained drug concentration. Mice treated with edaravone were sacrificed five minutes after the final intraperitoneal administration, when according to the previous report the maximum concentration was observed [Ito, 2008]. The plasma concentrations of EPI-589 and edaravone were measured by using liquid chromatography – tándem mass spectrometry (LC-MS/MS) technology, and expressed as mean ± standard deviation (SD; ng/mL).

The protein in plasma samples was precipitated by the addition of methanol containing internal standard (EPI-589-d5 for EPI-589 and phenytoin for edaravone). After centrifugation, an aliquot of the supernatant was diluted 1:2 with water. Aliquots of plasma extracts were analyzed by a Shimadzu LC-10 or 20A series high performance liquid chromatography system (Shimadzu Corporation) coupled with anAPI-4000 mass spectrometer (AB Sciex Pte. Ltd.) using a Unison UK-C18 HT column (3 μm, 75 mm × 2.0 mm i.d., Imtakt Corporation) for EPI-589 and Kinetex XB-C18 100A column (5 μm, 50 mm × 2.1 mm i.d., Phenomenex Inc.) for edaravone. The mobile phase consisted of 10 mM ammonium acetate (pH 4.0) as solvent A and methanol as solvent B. The gradient was performed at a total flow rate of 0.4 mL/min under the following conditions: for EPI-589, 0–0.3 min 10% (B), 0.3–2.5 min 10–90% (B), 2.5–3.7 min 90% (B), 3.7–3.71 min 90–10% (B), and 3.71–5.5 min 10% (B); for edaravone, 0–0.3 min 5% (B), 0.3–4.5 min 5–90% (B), 4.5–5.7 min 90% (B), 5.7– 5.71 min 90–5% (B), and 5.71–7.5 min 5% (B). The MS/MS transitions monitored in the positive ion mode were as follows: for EPI-589, m/z 266.2 → 203.2; for EPI-589-d5, m/z 271.2 → 207.2; for edaravone, m/z 174.9 → 133.1; for phenytoin, m/z 253.2 → 182.2.

### 2.8. Immunohistological analysis

The cervical spinal cord between C4 and C8 was embedded in paraffin, sectioned sagittally at 3 μm-thickness every 150 μm, and double-stained using the immunoenzyme method with anti-SMI-32 (BioLegend 801701, San Diego, CA, USA) and anti-MG-160 (Abcam ab103439, UK) antibodies used for the immunohistochemistry of motor neurons and the Golgi apparatus. SMI-32-positive (SMI-32+) neurons with diameters larger than 20 μm were defined as motor neurons (Hideyama et al., 2010). The number of motor neurons or those with MG-160 immunoreactivity (SMI-32+/MG-160+) in the cervical cord ventral horn or both were counted in 8 to 14 sections per mouse.

### 2.9. Measurement of pNfH level in plasma and 8-OHdG level in urine

The phosphorylated neurofilament H ELISA kit (BioVendor Laboratorni Medicina AS, Modrice, Czech Republic) was used for the pNfH assay. The measurement was once for each sample and according to the manufacturer’s instructions. Plasma samples were diluted 1:4 with dilution buffer. The level of pNfH of each sample was measured as an absorbance at 450 nm in a SpectraMax plate reader (Molecular Devices, LLC., San Jose, CA, USA). The 8-OHdG levels in urine collected from 7-week-old mice were measured once for each sample by using the 8-OHdG DNA Damage ELISA kit manufactured by Cell Biolabs, Inc. (San Diego, CA, USA), following the manufacturer’s protocol. Urine samples were diluted 1:10 with phosphate-buffered saline. The level of 8-OHdG of each sample was measured as an absorbance at 450 nm in a SpectraMax plate reader.

### 2.10. Tissue preparation and nuclear magnetic resonance (NMR) spectroscopy

The spinal cord tissue from the C4 to C6 region was collected in Eppendorf tubes and immediately frozen in liquid nitrogen. All tissues were stored at –80°C until the 1H NMR spectroscopy experiment. All NMR spectroscopy was performed and the data were recorded on an Agilent NMR system 400 MHz spectrometer (Agilent Techonology Inc., CA, USA) equipped with a Varian nano-probe at 4°C. The spinal cord tissue was dissected and packed into a 4 mm zirconium rotor (Agilent Techonology Inc., CA, USA) with 50 μL of D2O saline (0.9%) to provide magnetic field lock signal for the NMR spectrometer. The samples were spun at 2000 Hz to keep the rotation sidebands out of the acquisition window. One-dimensional (1D) 1H NMR spectra were acquired using a Carr-Purcell-Meiboom-Gill (CPMG) pulse sequence with 10,000 complex points, a 10,000 Hz spectral window, 2.0 sec presaturation delay, 1.0 sec acquisition time, 256 transients, 1.0 msec inter-pulse delay (τ), and 17 min 33 sec total experiment time.

The spectral data of the N-acetylaspartate (NAA) peak was analyzed with LCModel software (ver 6.3) and a set of basis spectra. As the calculation of absolute concentrations calibrated with precise sample volume and weight during rotation was difficult, the calculations of relative concentrations to the total creatine (creatine [Cr] plus phosphocreatine [PCr]) peaks were used for analysis. Data were excluded if the metabolite concentrations had an estimated standard deviation higher than 20% of the estimated concentration (Cramer-Rao Bounds [%SD]).

### 2.11. Hydroxyl radical scavenging activities

To evaluate hydroxyl radical scavenging activities, we used the Radical Catch assay kit (Hitachi, Ltd., Tokyo, Japan), which was a Fenton-type assay system, and followed the manufacturer’s protocol. In brief, using a 96-well microplate, a total volume of 120 μL including 5 μL of test article solution or ethanol for the vehicle, 50 μL of cobalt chloride solution, 50 μL of luminol solution, and 15 μL of distilled water was added to each well and mixed, and the microplate was put in a microplate reader set at 37°C for 5 min. Then 50 μL of hydrogen peroxide solution was immediately added to the reaction mixture, and the microplate was put back in a microplate reader and shaken for 2 sec. The luminescence of each reaction mixture was immediately measured every 12 sec for 3 min with the microplate reader set at 37°C. The assay performed in duplicate was repeated three times for EPI-589 and two times for DSR-303359 and edaravone. In each assay, hydroxyl radical scavenging activities (%) against vehicle were calculated from the mean values of luminescence of each drug concentration, and EC50 values in respective assays, where no luminescence was regarded as 100%, were determined as described in 2.13. Statistical analyses.

### 2.12. Cell viability assay

ALS patient derived fibroblast cells with fused in sarcoma (FUS) gene mutation (ND39027; Coriell Institute, NJ, USA) and SOD1 mutation (ND39026) were cultured in 384-well microplates at a density of 650 cells/well with assay media composed of 64% MEM-alpha, 25% Medium 199, 15% FBS, 1% penicillin-streptomycin, 10 mg/mL insulin, 10 μg/mL EGF and 10 μg/mL FGF. Five hours after the plating, 150 μM ferric citrate and test compounds were added to each well. On the next day, cultured cells were exposed for 2 days to 100 μM buthionine sulfoximine (BSO; Sigma-Aldrich, MO, USA), a glutathione synthase inhibitor, which leads to loss of cytoplasmic glutathione followed by oxidative stress and cell death (Wüllner et al., 1999; Sun et al., 2018). Viable cells were measured by the Calcein AM assay using the Cell Counting Kit-F (Dojindo Laboratories, Kumamoto, Japan). The assays performed in quadruplicate were repeated two times for both ALS fibroblasts. The mean values and standard error of the mean (SEM) of cell viabilities at each drug concentration were calculated, and EC50 values of EPI-589 and edaravone in each assay were determined, where cell viability in the BSO-free condition was regarded as 100%.

Cell viability was also examined in cultured mouse immortalized striatal STHdHQ7/Q7 cells under cystine depletion, which induces oxidative stress and consequent apoptotic cell death (Sun et al., 2018). Five hours after the plating of STHdHQ7/Q7 cells on 96-well microplates, the culture medium was changed to cystine-depleted medium containing 10% FBS, 1% penicillin-streptomycin, 4 mM glutamine, 1 mM pyruvate, and 30 mg/L methionine for 18 hours, and test compounds were added at the same time as the medium change. Then viable cells were measured by the Calcein AM assay using the Cell Counting Kit-F (Dojindo Laboratories). The assays of cystine deprivation-induced cell death performed in triplicate were repeated five times. The mean values and SEM of cell viabilities at each drug concentration were calculated, and EC50 values of EPI-589 and edaravone in each assay were determined, where cell viability in cystine-containing medium was regarded as 100%.

### 2.13. Statistical analyses

The mean, SD and SEM were calculated using Excel 2016 (Microsoft Corporation). Statistical analyses were performed using SAS (SAS Institute Inc., version 9.2) and Stat Preclinica (Takumi Information Technology, Inc., version 1.2). Homogeneity of the variances between groups in the wobbler mice behavioral study were evaluated by Bartlett’s test. Significance of the differences among the values of rotarod walking time and forelimb deformity score at each time point in wobbler mice were evaluated by two-way repeated measures analysis of variance (ANOVA) followed by Dunnett’s post hoc test. Differences between groups with respect to SMI-32 positive neurons with diameters larger than 20 μm per slice in the histological study and biomarkers such as pNfH level in plasma, 8-OHdG level in urine, and spinal NAA level measured by MRS were evaluated by the Student t-test. Two-sided p-values < 0.05 were considered to be statistically significant. EC50 values and 95% confidence interval (95% CI) were calculated using logistic regression analysis.

## 3. Results

### 3.1. Hydroxyl radical scavenging activities of EPI-589, DSR-303359, and edaravone

The hydroxyl radical scavenging activities of EPI-589, DSR-303359, and edaravone were assessed in cell free supernatants using a Fenton’s reaction-luminol chemiluminescence system for detection of hydroxyl radicals (Fig. 2). A free radical scavenger edaravone demonstrated dose-dependent hydroxyl radical scavenging activity with mean EC50 of 2.22 μM (2.53 and 1.91 μM in two independent assays), which is consistent with the previous study (Watanabe et al., 2008). EPI-589 had no radical scavenging activity at concentrations at least up to 30 μM in three assays, whereas DSR-303359, a reduced form of EPI-589, showed potent and dose-dependent scavenging activity with EC50 of 1.84 μM (1.11 and 2.56 μM in two independent assays), which was almost equivalent to that of edaravone.

**Fig. 2.**
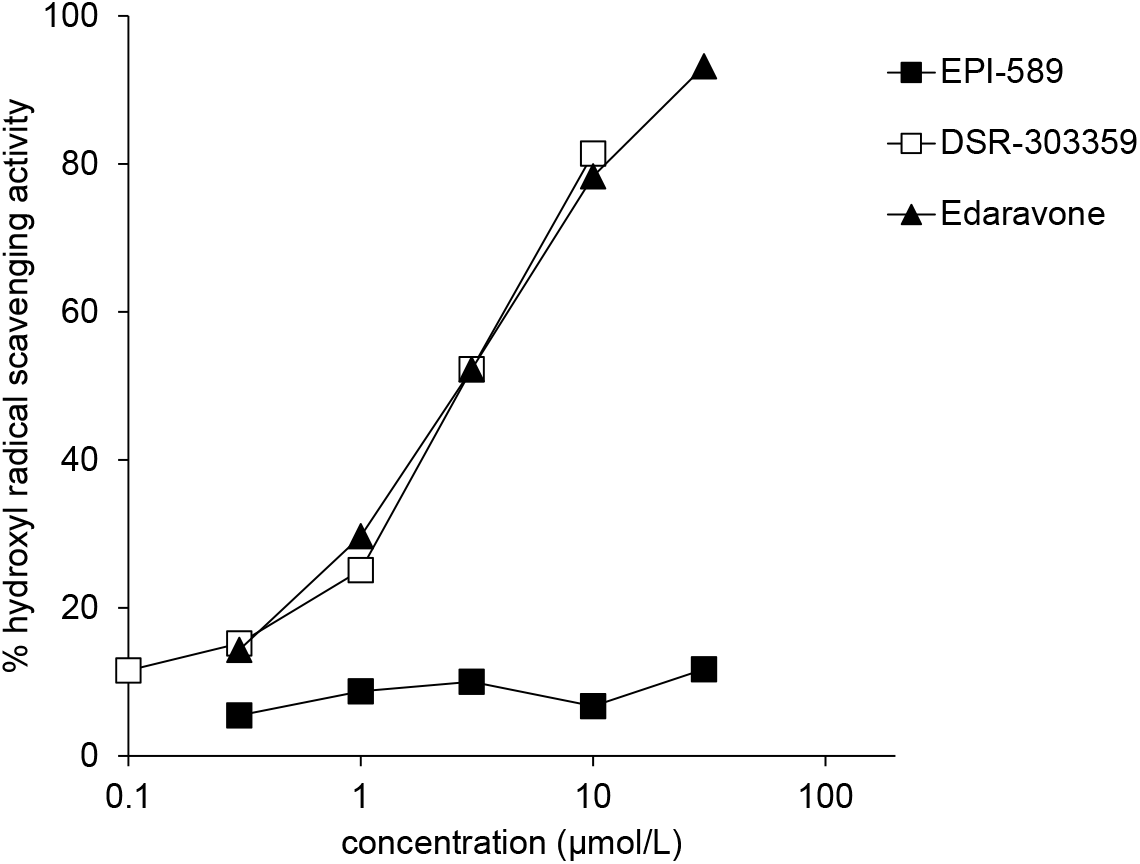
Hydroxyl radical scavenging activities under cell-free conditions. Representative data are shown the hydroxyl radical scavenging activities of EPI-589, DSR-303359, and edaravone. In this assay edaravone showed scavenging activity against hydroxyl radicals generated by the Fenton reaction. EPI-589 was not effective, whereas DSR-303359, a reduced form of EPI-589, had potent hydroxyl radical scavenging activity. Each assay was performed in duplicate and the representative data are expressed as the mean value of two wells.

### 3.2. Effect of EPI-589 and edaravone on cell viability in BSO-treated and ferric citrate-treated ALS fibroblast cells

Fibroblasts from FUS or SOD1 mutant ALS patient died after exposure to BSO and ferric citrate for 2 days as described in the previous study (Wüllner et al., 1999; Sun et al., 2018). EPI-589 strongly attenuated cell death caused by BSO and ferric citrate with mean EC50 of 0.187 μM (0.140 and 0.235 μM in two assays) in FUS-ALS fibroblasts and 0.114 μM (0.169 and 0.058 μM in two assays) in SOD1-ALS fibroblasts, respectively (Fig. 3a). On the other hand, edaravone showed only a partial protective effect in this cellular model; the EC50 of edaravone was estimated to be higher than 250 μM.

**Fig. 3.**
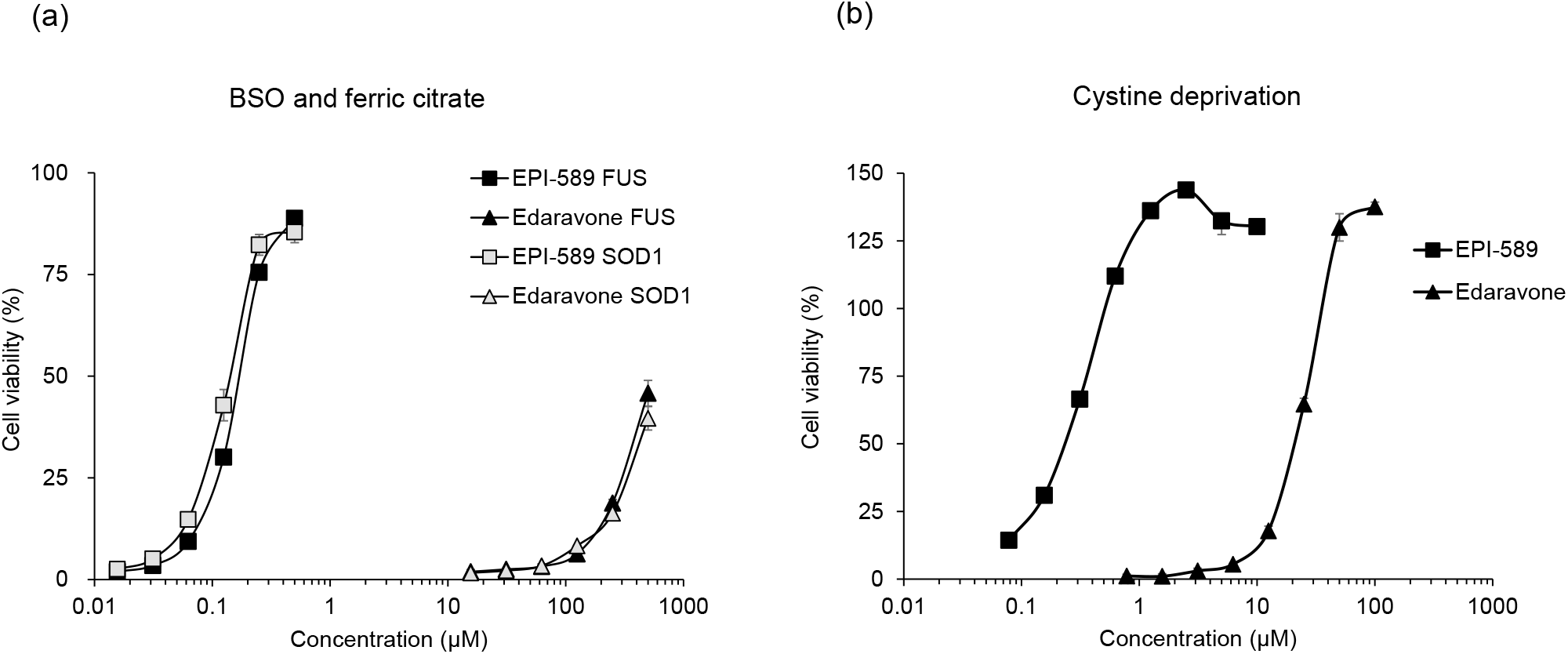
Protective effects of EPI-589 and edaravone against glutathione depletion and cystine depletion in culture cells. Representative data are shown from the assays evaluating the effects of EPI-589 and edaravone on cell viability in BSO-treated and ferric citrate-treated ALS fibroblast cells (a), and cystine-depleted mouse STHdHQ7/Q7 cells (b). EPI-589 had much more potent protective effects than edaravone on cells under both oxidative stress conditions. Data are expressed as mean ± SEM.

### 3.3. Effect of EPI-589 and edaravone on cell viability in cystine-depleted mouse STHdHQ7/Q7 cells

Mouse immortalized striatal STHdHQ7/Q7 cells that were cultured in cystine-free medium for 18 hours showed cell death, which is consistent with observation in the previous study (Sun et al., 2018). EPI-589 strongly attenuated the cell death of mouse immortalized striatal STHdHQ7/Q7 cells caused by cystine deprivation (Fig. 3b) with mean EC50 of 0.45 μM (95% CI, 0.26 – 0.65 μM) from five independent assays. Edaravone also showed a protective effect against cystine depletion-induced cell death; however, the EC50 of edaravone was 24.09 μM (95% CI, 21.23 – 26.94 μM), which is much higher than that of EPI-589.

### 3.4. Effect of EPI-589 and edaravone on motor function and clinical symptoms in wobbler mice (behavioral study)

The wobbler mice in the control group showed progressive reduction of rotarod walking time throughout the experimental period, as shown in Fig. 4a. The wobbler mice in the EPI-589 group showed an almost constant rotarod walking time of 300 sec up to day 18 of treatment and had significantly greater walking time from day 19 to 33 of treatment, compared with that in the control group (p = 0.0001, 0.0037, 0.0018 and 0.0014 at days 19, 26, 30 and 33, and p < 0.0001 at day 23; two-way repeated measures ANOVA followed by Dunnett’s test). On the other hand, the mice in the edaravone group showed greater walking time that was only temporary and moderate at day 18 of treatment (p = 0.0345 vs. that in the control group), and did not show any difference in walking time at day 21 of treatment or later. The wobbler mice in the control group also showed progressive worsening of forelimb deformity score throughout the experimental period (Fig. 4b). However, in the EPI-589 group, they showed significantly lower forelimb deformity scores at day 2 (p = 0.0385 vs. that in the control group) and several time points after day 12 of treatment (p = 0.0064, 0.0005, 0.0007 and 0.0013 at days 12, 19, 23, 30, and p < 0.0001 at day 33 vs. that in the control group), whereas in the edaravone group, they did not show any difference in forelimb deformity score throughout the experimental period, compared with that in the control group.

**Fig. 4.**
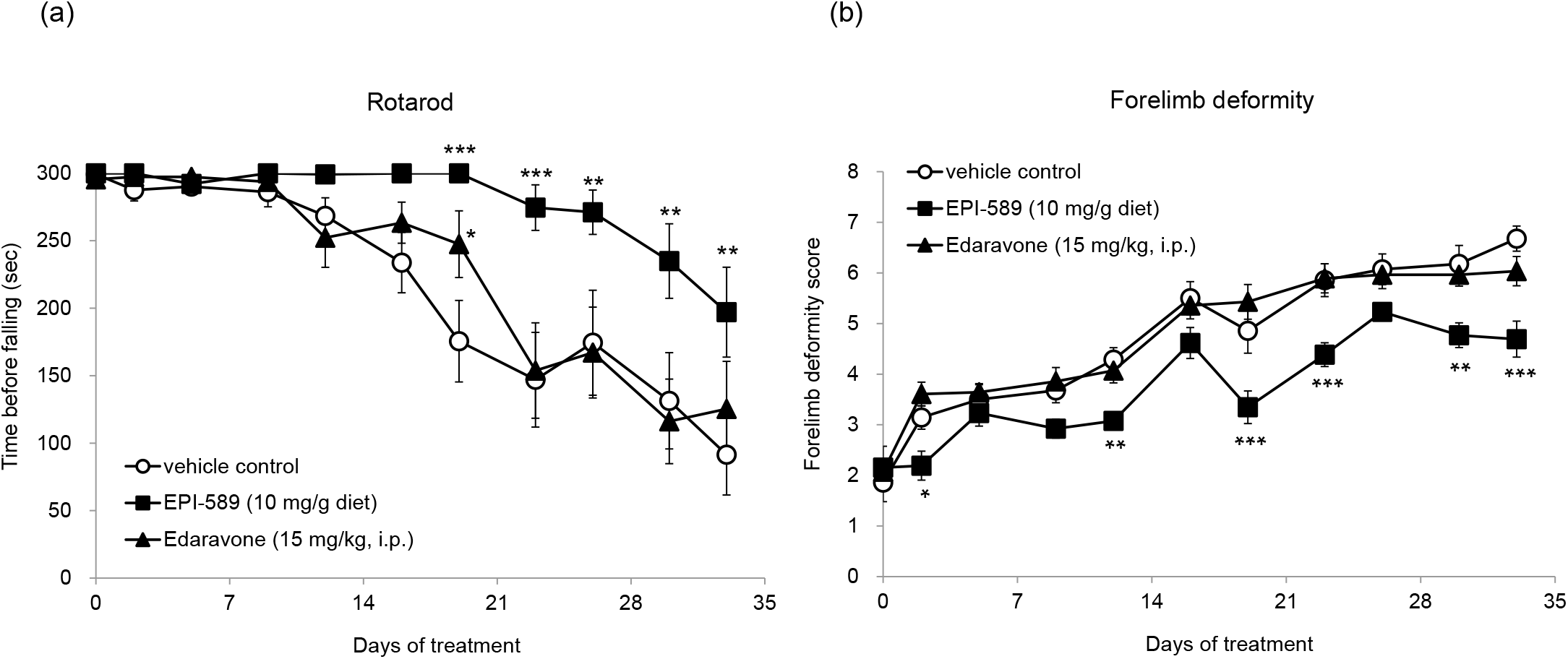
Effect of EPI-589 and edaravone on motor deficit and forelimb deformity in wobbler mice. EPI-589 delayed the deterioration of rotarod walking performance (a) and development of forelimb deformity (b) in wobbler mice. The number of animals in vehicle control, EPI-589 and edaravone groups were 14, 13, and 14, respectively. Data are expressed as mean ± SEM. *p < 0.05, **p < 0.01, ***p < 0.001 vs vehicle control group in two-way repeated measures ANOVA followed by Dunnett’s post hoc test.

There were significant differences from the control group in body weight in the EPI-589 group at day 9 and in the edaravone group at days 12, 16, and 19, however these differences were only temporary (Supplementary Fig. 1). Moreover, efficacy of EPI-589 was observed in both gender in rotarod and forelimb deformity test (Supplementary Fig. 2). The plasma concentration of EPI-589 at the end of the treatment was 901 ± 781 ng/mL (3.40 ± 2.94 μM, mean ± SD), which was close to the lower end of the concentration as observed in the preliminary study (see Materials and Methods). The plasma concentration of edaravone at the end of the treatment was 5761 ± 1541 ng/mL (33.07 ± 8.85 μM).

### 3.5. Effects of EPI-589 on pathophysiological changes in wobbler mice (biomarker study 1 and 2)

The histopathological analysis in biomarker study 1 demonstrated that nearly all of the motor neurons (SMI-32+ neurons with diameters larger than 20 μm) in the cervical cord ventral horn were immunopositive also for MG-160 in wild type mice at 6 weeks of age, whereas a substantial number of motor neurons were not immunoreactive for MG-160 in age-matched wobbler mice in the control diet group (Fig. 5a, b). Accordingly, the number of SMI-32 and MG-160 double-positive (SMI-32+/MG-160+) motor neurons was significantly lower in the wobbler mice than in wild type littermates (Fig. 5b, n = 4, p = 0.0049 in the Student t-test), while there were no significant differences in the total number of motor neurons between the two groups. Interestingly, the number of SMI-32+/MG-160+ motor neurons was significantly greater in the group of wobbler mice fed EPI-589 from 4 to 6 weeks than in those fed the control diet over the same period (Fig. 5b, n = 4, p = 0.0232 in the Student t-test).

**Fig. 5.**
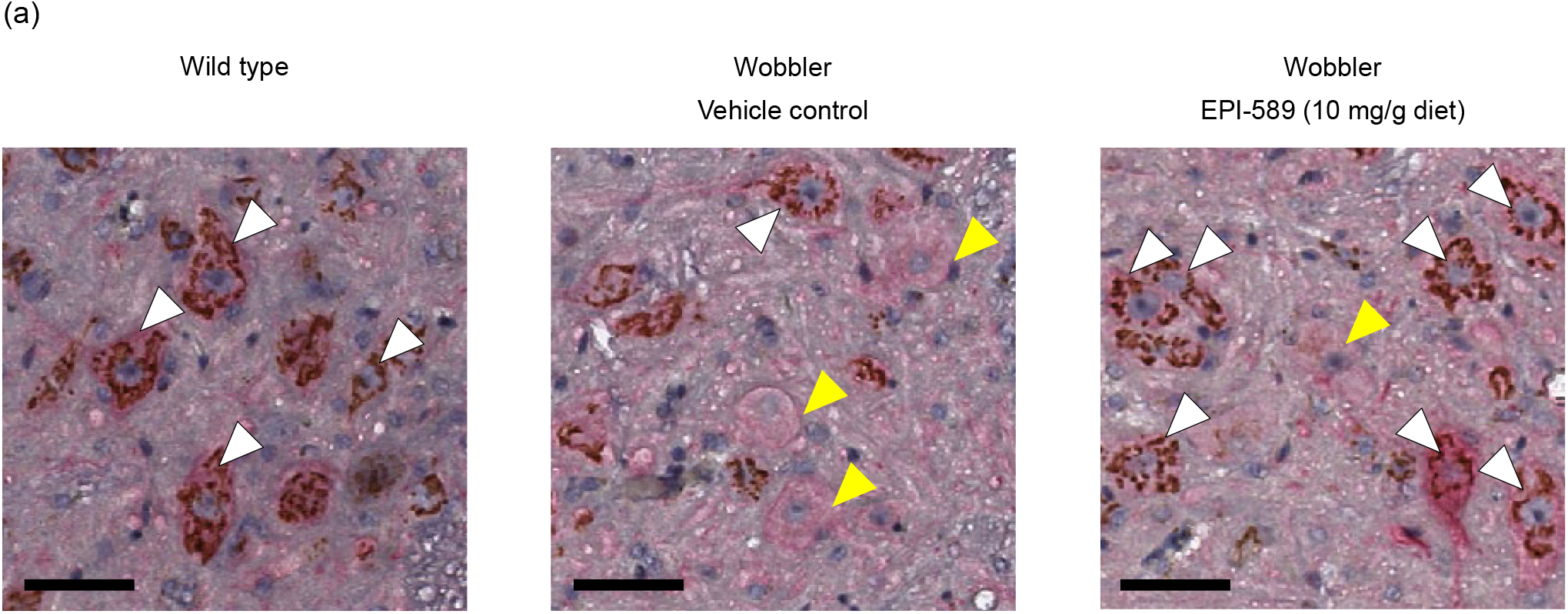
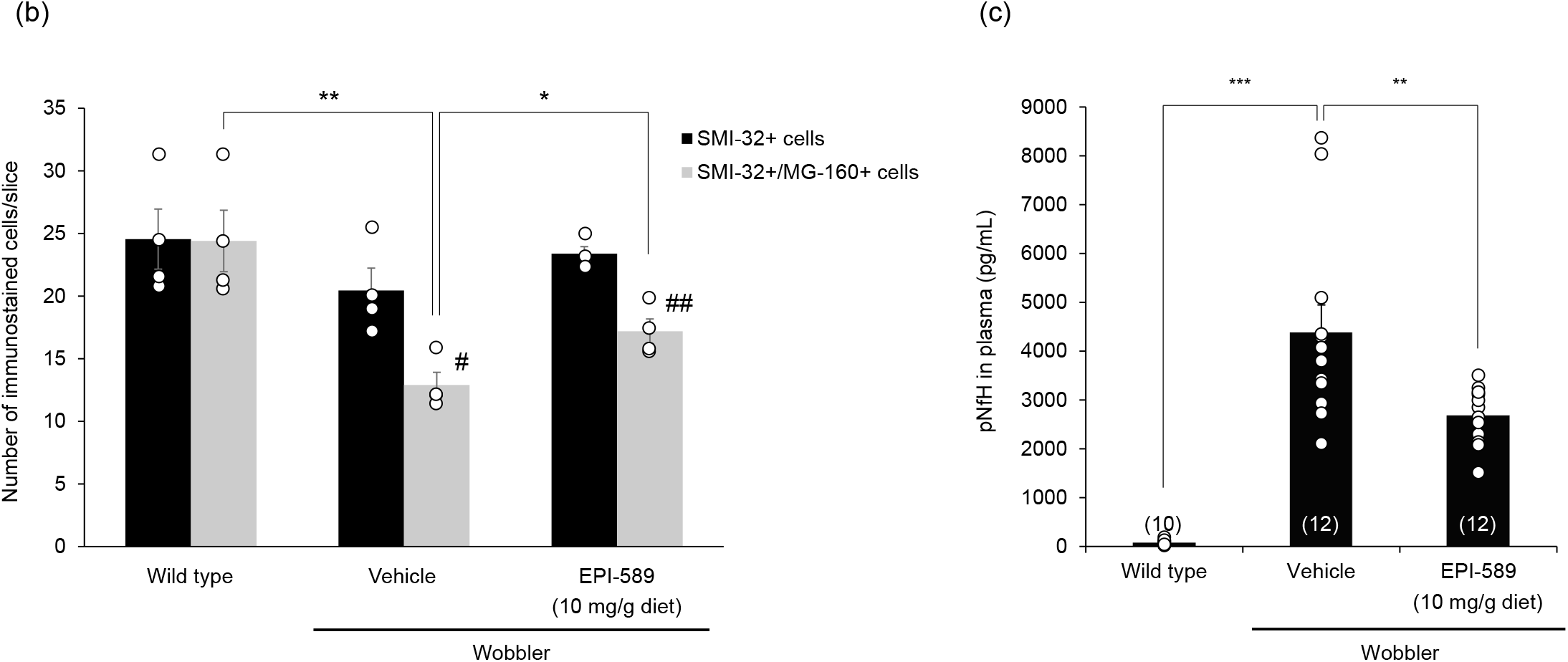
Effect of EPI-589 (10 mg/g of diet) on pathological changes of the cervical cord motor neurons (a, b), and plasma pNfH (c) in the wobbler mouse biomarker study 1. (a) A representative image of motor neurons double immunostained with anti-SMI-32 and anti-MG-160 antibodies in wild type mouse, wobbler mouse fed the normal diet, and wobbler mouse fed the EPI-589-containing diet. White arrowheads indicate SMI-32 and MG-160 double-positive (SMI-32+/MG-160+) motor neurons, and yellow arrowheads indicate SMI-32+ motor neurons without MG-160 immunostaining. The motor neurons without MG-160 immunostaining are in the ventral horn of cervical spinal cord of wobbler mice. Scale bars, 50 µm. (b) The numbers of total SMI-32+ motor neurons (MNs) and SMI-32+/MG-160+ MNs per slice. Data are expressed as mean ± SEM (n = 4 in each group). *p < 0.05, **p < 0.01 between groups, #p < 0.05, ##p < 0.01 between the numbers of SMI-32+ MNs and SMI-32+/MG-160+ MNs in each group, in the Student t-test. (c) Plasma pNfH was measured by ELISA. EPI-589 significantly prevented the elevation of the pNfH level in plasma collected from 6-week-old wobbler mice. Numbers in parentheses show the number of mice examined. Data are expressed as mean ± SEM. **p < 0.01, ***p < 0.001 in the Student t-test.

The analysis of plasma pNfH level in biomarker study 1 demonstrated that it was significantly greater in the wobbler mice fed the control diet at 6 weeks of age than in age-matched wild type littermates (Fig. 5c, n = 10 – 12, p < 0.0001 in the Student t-test). The plasma pNfH level was significantly lower in the wobbler mice fed dietary EPI-589 than in those fed the control diet (Fig. 5c, n = 12, p = 0.0085 in the Student t-test).

The urinary 8-OHdG level examined in biomarker study 2 demonstrated that it was significantly greater in the wobbler mice fed the control diet at 7 weeks of age than in age-matched, wild type littermates (Fig. 6a, n = 13 – 14, p = 0.0104 in the Student t-test). On the other hand, the plasma 8-OHdG level was significantly lower in the wobbler mice fed dietary EPI-589 than in those fed the control diet (Fig. 6a, n = 12 – 14, p = 0.0042 in the Student t-test). In another set of animals used to compare the ratio of NAA to creatine (Cr) plus phosphocreatine (PCr) (NAA/[Cr+PCr]) in the cervical cord tissues between the wobbler and age-matched wild type mice at 8 weeks of age, the NAA/(Cr+PCr) ratio was significantly lower in the wobbler mice than in wild type mice (Fig. 6b, n = 6, p = 0.0260 in the Student t-test) and significantly greater in the wobbler mice fed dietary EPI-589 than in those fed the control diet (Fig. 6b, n = 6, p = 0.0032 in the Student t-test).

**Fig. 6.**
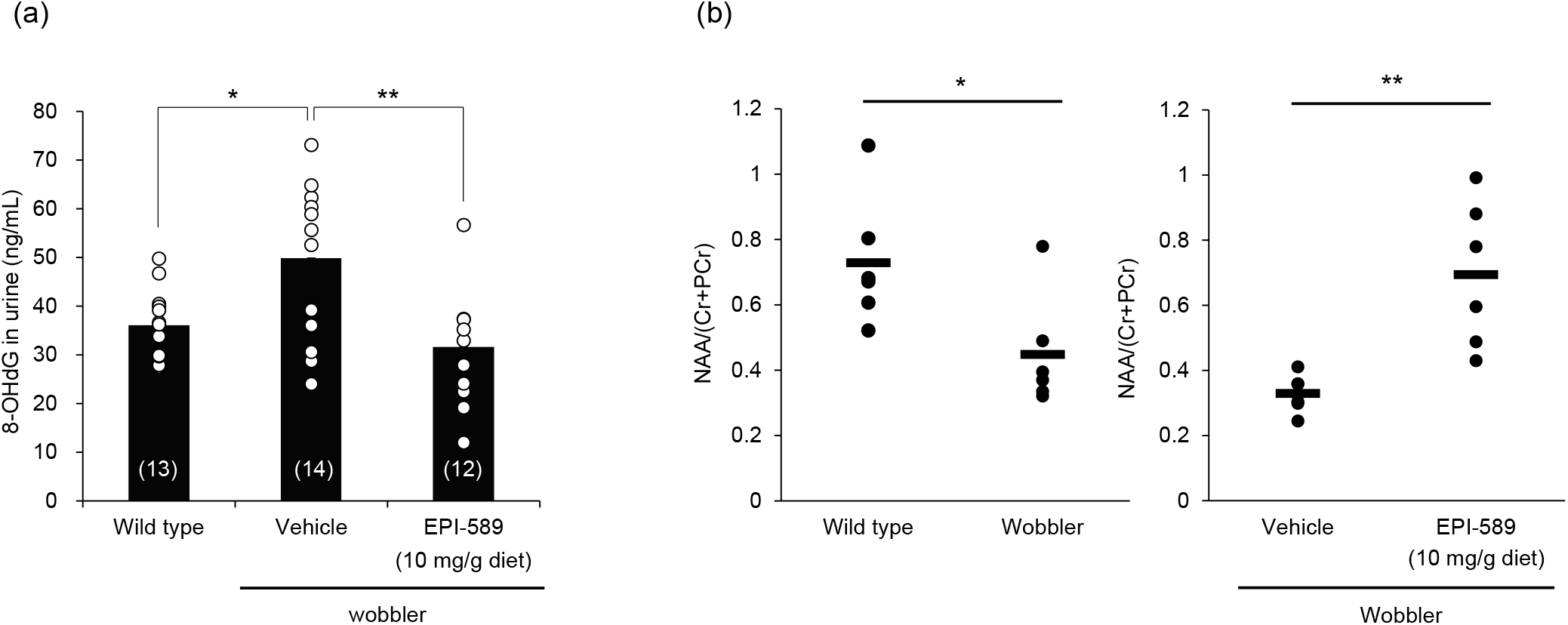
Urinary 8-hydroxy-2’-deoxyguanosine (8-OHdG) (a) and cervical cord N-acetylaspartate (NAA) (b) were examined in biomarker study 2. (a) 8-OHdG level in urine was measured by ELISA. EPI-589 (10 mg/g diet) reduced the level of 8-OHdG in urine. Numbers in parentheses show the number of mice examined. Data are expressed as mean ± SEM. *p < 0.05, **p < 0.01, in the Student t-test. (b) Cervical cord NAA was measured by nuclear magnetic resonance spectrometry (NMRS). The ratio of NAA to creatine (Cr) plus phosphocreatine (PCr) (NAA/[Cr+PCr]) in the cervical cord tissues of wobbler mice at 8 weeks of age was significantly lower than that of age-matched wild type mice. EPI-589 significantly inhibited the decrease of the NAA level normalized to creatine (Cr) plus phosphocreatine (PCr) levels in cervical cord tissues of wobbler mice at 8 weeks of age. Mean and individual data are indicated by a bar and circles, respectively (n = 6 in each group). *p < 0.05, **p < 0.01 in the Student t-test.

## 4. Discussion

EPI-589 has been designed as a cell membrane-penetrable, small molecule with benzoquinone structure. Various benzoquinone-containing compounds are well known to occur naturally in plants, animals, and bacteria, where they play various physiological roles such as bioenergetic transport, oxidative phosphorylation, and electron transfer (Dandawate et al., 2010). Among them, the reduced form of benzoquinone-containing compounds with antioxidant activity such as ubiquinol has been extensively investigated. Ubiquinol is enriched in the living body to defend against oxidative stress efficiently through reductive enzymatic mechanisms that convert it from endogenously synthesized ubiquinone (Bentinger et al., 2007). Likewise, EPI-589 is the oxidized form, which is supposedly easily reduced in the living body into an antioxidant form, as exemplified above by the reduction of ubiquinone to ubiquinol. The present study firstly tested this hypothesis. Actually, *in vitro* experiments in the present study demonstrated that in cell-free supernatants, only the reduced form (DSR-303359) but not the oxidized form (EPI-589) had hydroxyl radical trapping activity, whereas in cell culture, EPI-589 itself turned out to have highly potent cell-protective activity in well-established models of oxidative stress such as the one induced by glutathione depletion, supporting well our hypothesis.

Surprisingly, the present study found that EPI-589 could be exceedingly more protective of cell viability against oxidative stress than the free-radical scavenger edaravone (more than 1300 times more protective with a difference in EC50 of nearly 50 times in ALS fibroblasts and mouse STHdHQ7/Q7 cells, respectively) under the cell culture conditions used. However, in cell-free supernatants, EPI-589 itself was not efficacious and the reduced form of EPI-589 was nearly equivalent to edaravone in its ability to scavenge hydroxyl radical. The different molecular mechanisms that these compounds use to defend against oxidative stress may explain the striking differences in antioxidant potencies under cell culture conditions. For example, as described above, EPI-589 is a sort of a catalytic compound that can be recycled to the reduced form in the cells by redox enzymes, whereas edaravone is a free radical scavenger that is metabolized to the oxidized form by trapping free radicals, and is supposedly not recycled to the reduced form (Watanabe et al., 2004). Therefore, EPI-589 can eliminate free radicals more efficiently with the support of redox enzymes under cell culture conditions than under cell free conditions, whereas edaravone can act similarly under both of these conditions (Watanabe et al., 2004). In contrast to edaravone, EPI-589 may function as a redox active benzoquinone-containing compound like ubiquinone to facilitate mitochondrial electron transport and prevent mitochondrial energy depletion and subsequent apoptosis, which has been previously shown to occur following reactive oxygen species generation by GSH depletion with BSO (Merad-Boudia et al., 1998). However, these intracellular mechanisms of EPI-589 in comparison with that of edaravone were not addressed by the present study and should be clarified in detail by further experiments.

The present study successfully replicated the previous finding that daily intraperitoneal administration of edaravone delayed symptomatic progression of motor neuron disease in wobbler mice, although the efficacy of edaravone in the present study was only transient and not seen in the scoring offorelimb deformity, in contrast to the previous study (Ikeda and Iwasaki, 2015). It is likely that the differences in efficacy may be due to differences in the study design between studies, i.e., in this study, treatment began a little later (4 weeks vs 3 – 4 weeks), dose level was greater (15 vs 10 mg/kg, i.p.), housing conditions were different with a normal littermate placed in each cage as a foster mouse, and so on. Regarding the dosage, we attempted to use the same dose (daily 15 mg/kg, i.p.) in this experiment as previously done in experiments using the SOD1-transgenic mouse model, however, the plasma concentration (5761 ± 1541 ng/mL, 33.07 ± 8.85 μM) at 5 min after the final dosage in wobbler mice was unexpectedly greater than that estimated from the previous study in SOD1 mice (1286 ng/mL or 1.5 times that of 10 mg/kg dose; Ito et al., 2008), indicating a possible difference in the pharmacokinetics of edaravone between these mouse models. It has been also reported that the maximum concentration of edaravone in human plasma at the clinical dose used for treating ALS in patients was estimated to be approximately 1000 ng/mL (Nakamaru et al., 2017), which appears to be less one fifth the concentration observed in the present study. Thus, the dosage of edaravone used in this study is considered to have achieved a greater plasma concentration than the dosages used in previous mouse model and ALS studies (Ikeda and Iwasaki, 2015; Ito et al., 2008; Edaravone (MCI-186) ALS 19 Study Group, 2017). Nevertheless, edaravone in this study was found to be well-tolerated in wobbler mice and considered to be appropriate to use for evaluating the efficacy in this animal model. Replication of the edaravone’s efficacy for motor symptoms in wobbler mice support the notion that free radical generation and oxidative stress play a significant role in this model of motor neuron disease (Santoro et al., 2004; Moser et al., 2013).

Very interestingly, the present study found that dietary EPI-589 very robustly delays the symptomatic progression of motor neuron disease in the wobbler mouse, in contrast to edaravone. It is considered that differences in both the pharmacokinetics and pharmacodynamics between these compounds likely contribute to the difference in efficacy between these compounds in the wobbler mouse. From the point of view of pharmacokinetics, we also observed that, though the plasma concentration of EPI-589 in wobbler mice was 901 ng/mL (3.40 μM) at the end of treatment for 5 weeks in this study, it could range between 737 and 2970 ng/mL (2.78 and 11.19 μM) in the preliminary study to measure the plasma concentration every 4 hours for 4 days after dietary dosing of EPI-589. Notably, the maximum exposure level may be less after dosing with dietary EPI-589 than with edaravone (i.p.), whereas the minimum exposure level may be greater after dosing with dietary EPI-589 than with edaravone (i.p.), and the minimum exposure level to edaravone should be negligible based on edaravone’s pharmacokinetics as described previously (Nakamaru et al., 2017). Therefore, it is possibly important to maintain for a certain period of time the appropriate exposure level in a concentration range that achieves robust efficacy as seen with dietary EPI-589 but not edaravone (i.p.) in this study. Pharmacodynamically speaking, since the present study discovered that EPI-589 could much more potently protect cells against oxidative stress than edaravoneunder some culture conditions, this potency difference due to different mechanisms of action (as discussed above) may contribute to the greater efficacy of EPI-589. Actually, the plasma concentration of EPI-589 in the wobbler mouse (3.40 μM) was greater than the *in vitro* cell-protective EC50 (0.114 to 0.45 μM), whereas that of edaravone *in vivo* (33.07 μM) appears to be equivalent to or less than its *in vitro* EC50 (24.09 μM or higher than 250 μM), demonstrating that *in vivo* exposure to the former but not the latter might be robustly efficacious enough *in vivo* to prevent or delay progression of motor neuron deterioration of wobbler mice. However, more detailed analysis of the relationship between the pharmacokinetics and pharmacodynamics for EPI-589 is apparently required to clarify the mechanisms underlying the efficacy of EPI-589 in the wobbler mouse model. Nevertheless, the present study revealed that protective therapy for oxidative stress could have robust efficacy for motor neuron disease, based on the suspected appropriate mechanism and potency of antioxidant action as well as the appropriate pharmacokinetics of EPI-589 in this study.

We explored widely the potential usefulness of various biological markers in this study to detect the pathophysiological changes in the wobbler mouse model of motor neuron disease, and tested the effects of EPI-589 to ameliorate the disease-related changes by utilizing such markers. Greater urinary level of 8-OHdG in wobbler mice fed the control diet than in those fed dietary EPI-589 or in normal littermates is consistent with the idea that oxidative stress occurring in wobbler mice (Santoro et al., 2004; Moser et al., 2013) may involve the oxidative damage of DNA, and EPI-589 may protect against such oxidative events while delaying the symptomatic progression of the disease. It is also reasonable to note the co-presentation of a greater level of plasma pNfH, lower number of pNfH positive motor neurons, and lower NAA/(Cr+PCr) ratio in the cervical cord in wobbler mice fed the control diet than in normal littermates until 8 weeks of age, which consistently support previous findings that cervical cord motor neurons are significantly degenerated by this age in wobbler mice (Mitsumoto and Bradley, 1982; Pollin et al., 1990). The pathological changes of these markers were observed to be diminished to various degrees by treatment with dietary EPI-589 in wobbler mice, suggesting that EPI-589 may delay the pathological progression as well as the symptomatic progression of motor neuron disease in this mouse model. Importantly, all of these markers such as 8-OHdG level (Bogdanov et al., 2000), pNfH level (Li et al., 2016), and NAA/(Cr+PCr) ratio (Carew et al., 2011) are measurable in ALS patients, suggesting the potential usefulness of these markers to evaluate the effects of EPI-589 clinically in ALS. In this regard, it is crucial to conduct an additional study in wobbler mice to investigate in more detail the natural history of these markers as the disease progresses to understand better the meaning of each biomarker in the disease’s pathophysiology and the therapeutic value of a drug candidate such as EPI-589.

Fragmentation of the Golgi apparatus in ALS was initially discovered by Gonatas’s group in 1990 (Mourelatos et al., 1990) with the use of an organelle-specific antibody MG-160 and then confirmed in sporadic, familial, and juvenile forms of ALS as well as SOD1 transgenic mice (Fujita and Okamoto, 2005). Golgi apparatus fragmentation is suggested to be a common pathological event in motor neuron disease that may occur at a relatively early phase prior to cytoskeletal lesions and neurodegeneration (Sundaramoorthy et al., 2015). Motor neuron disease in wobbler mice is known to be caused by a point mutation of the gene for Vps54, which is a component of the Golgi-associated retrograde protein (GARP) complex, with the complex serving as a vesicle-tethering factor for both early and late endosomes in the trans Golgi network (Schmitt-John et al., 2005; Moser et al., 2013). Pathologically, enlarged endosome vacuolization has been also confirmed as the earliest event in cervical cord motor neurons, occurring as early as 2 weeks of age (Mitsumoto and Bradley, 1982). Thus, it is reasonable to suspect that the fragmentation of the Golgi apparatus as detected by loss of MG-160 immunoreactivity may be also the case in the wobbler mouse model. Actually, we have successfully found in this study that the MG-160 immunoreactivity of a subpopulation of cervical cord motor neurons (approximately 27% of the total) was diminished in wobbler mice by 6 weeks of age. The Vps54 gene mutation has not been identified in ALS patients (Corrado et al., 2011), which may be one of limitations in the present study using wobbler mouse as a motor neuron disease model, however, the previous and present studies consistently suggest that the pathophysiological process in the wobbler mouse may resemble that of ALS, at least partly, and include the vesicle trafficking defect (Sundaramoorthy et al., 2015). Very interestingly, dietary EPI-589 treatment led to the increase of SMI-32+/MG-160+ motor neurons in wobbler mice in this study, suggesting that EPI-589 may ameliorate the pathological changes in the Golgi apparatus, which could be controlled by upstream mechanisms underlying the preservation of motor neuron structure and function observed with this drug treatment.

In conclusion, the present study firstly demonstrated that EPI-589 is a highly potent, redox-active neuroprotectant with robust effects that delay the symptomatic and pathophysiological progression of motor neuron disease in the wobbler mouse, possibly by counteracting the vesicle trafficking defect and oxidative stress causing motor neuron degeneration. The study also identifies the potential biomarker candidates, which may be useful for translational validation of the effects of EPI-589 on pathophysiological events in ALS. These findings strongly encourage further exploration of the therapeutic potential of EPI-589 in ALS.

## Author contributions

Y.M., K.S., T.N., M.Y., and T.I. provided substantial contributions to the study design. Y.F., N.T., and F.I. were involved in the data collection and analysis. Y.M. and T.I wrote the manuscript. All authors were involved in data interpretation. All authors discussed and agreed on the content of the final manuscript.

## Acknowledgments

We would like to thank Kan Miyoshi, Masakazu Isobe, Maiko Kitaichi, Natsuko Goto, and Yuka Yamana, members of Sumitomo Dainippon Pharma Co, Ltd., for their contribution to data collection.

---

**Supplementary Fig. 1.**
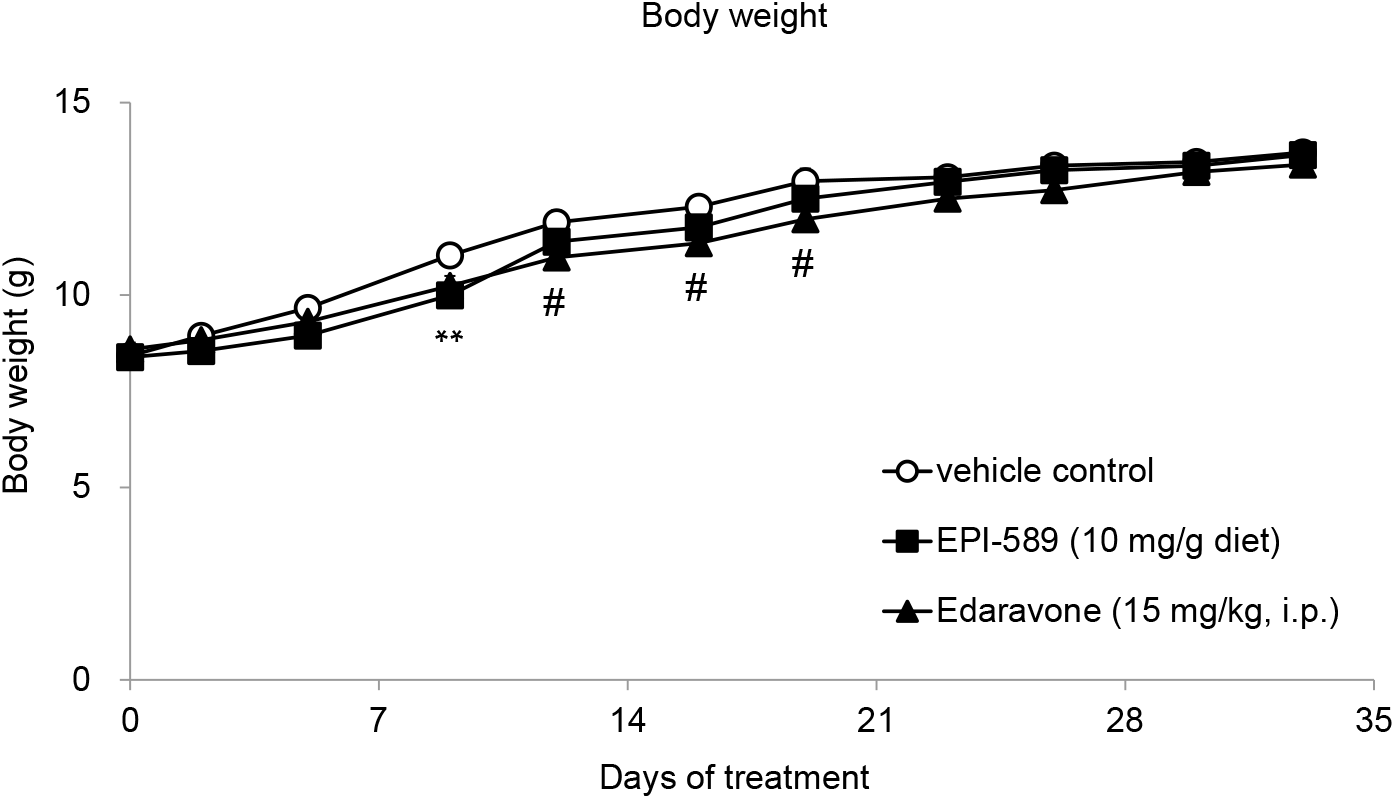
Effect of EPI-589 and edaravone on body weight in wobbler mice. The number of animals in vehicle control, EPI-589 and edaravone groups were 14, 13, and 14, respectively. Data are expressed as mean ± SEM. **p < 0.01 between vehicle control and EPI-589, #p < 0.05 between vehicle control and edaravone, in two-way repeated measures ANOVA followed by Dunnett’s post hoc test.

**Supplementary Fig. 2.**
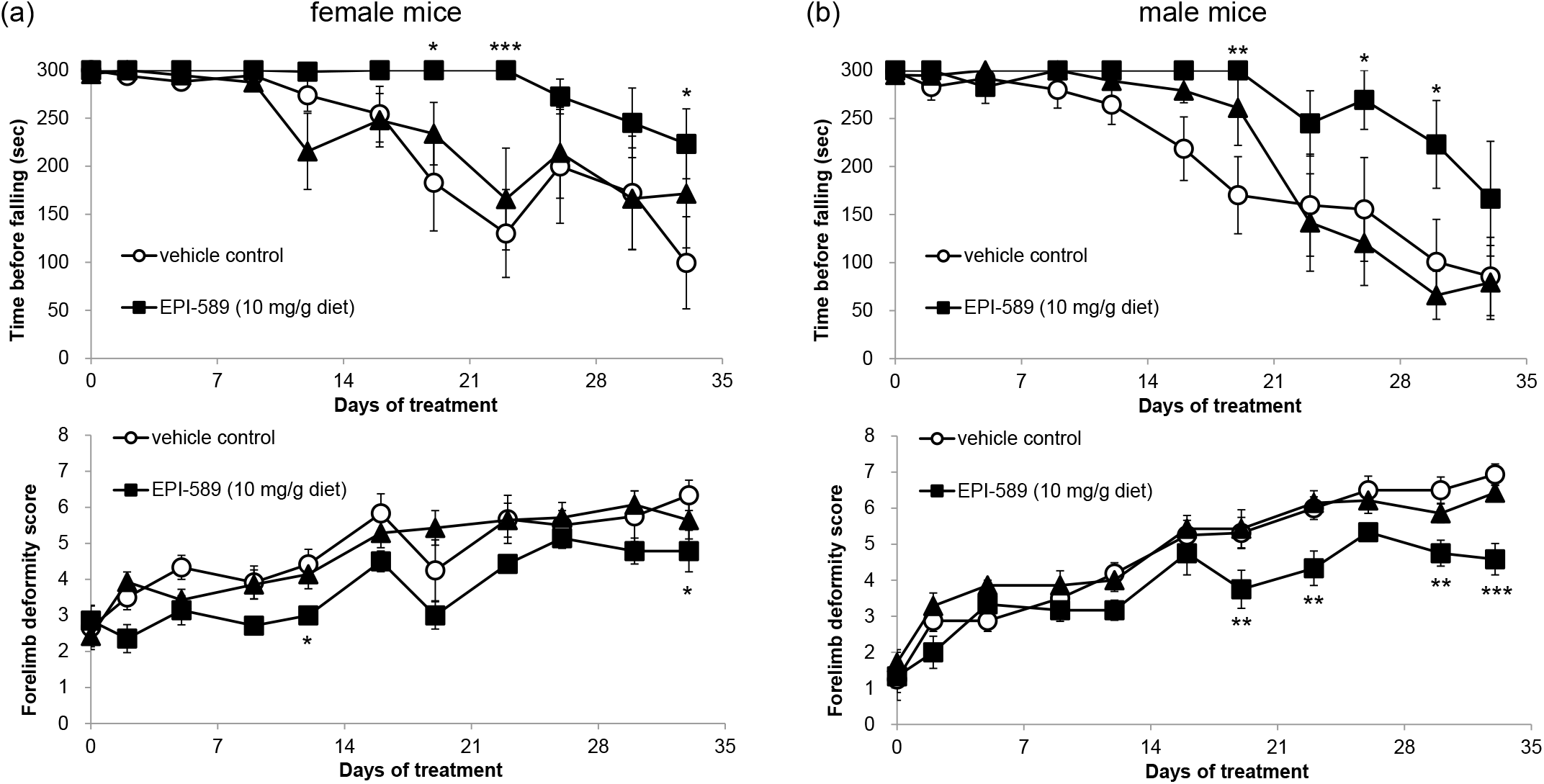
Effect of EPI-589 and edaravone on motor deficit and forelimb deformity in female (a) and male (b) wobbler mice. EPI-589 delayed the deterioration of rotarod walking performance and development of forelimb deformity in both gender in this study. The number of animals in vehicle control, EPI-589 and edaravone groups were 6, 7, and 7 in female mice, and 8, 6, and 7 in male mice, respectively. Data are expressed as mean ± SEM. *p < 0.05, **p < 0.01, ***p < 0.001 vs vehicle control group in two-way repeated measures ANOVA followed by Dunnett’s post hoc test.

